# A Single Structure-Derived Computational Metric Predicts High-Affinity Antibody Selection Against a Malaria Antigen

**DOI:** 10.1101/2025.09.30.679617

**Authors:** Gordon A. Dale, Bingxian Chen, Ja-Hyun Koo, Martina Wecera, Sven Kratochvil, Facundo D. Batista

**Affiliations:** Batista Lab, The Ragon Institute of Mass General Brigham, MIT, and Harvard, Cambridge, MA 02139, USA; Department of Biology, Massachusetts Institute of Technology, Cambridge, MA 02139, United States; Department of Immunology, Harvard Medical School, Boston, MA 02115, USA

**Author notes:** Liu Lab, The Ragon Institute of Mass General Brigham, MIT, and Harvard, Cambridge, MA 02139, USA.

## Abstract

There is an increasing need for improved malaria antibodies that can be used in passive immunization strategies to reduce the burden of malaria in endemic regions. Despite considerable progress, the identification or development of variants that meet stringent performance requirements remains a challenge. A key strategy has been the improvement of prototypic antibodies targeting the repeat antigens on *Plasmodium falciparum* circumsporozoite protein (PfCSP). In this work, we derive a computational metric from predicted protein structures that efficiently captures affinity information of antibody variants of the PfCSP-targeting antibody, CIS43. We then use this metric to rapidly explore sequence space as large as >3×10^47 variants using principles of the germinal center, deriving new high-affinity CIS43 variants from the method. We further extend this framework to generate high-affinity variants of an unrelated PfCSP-targeting antibody, L9, by maturing both homotypic and antigen-binding interactions, which demonstrates substantial flexibility of the approach. Taken together, we show that coupling micro-evolutionarily selected mutations to in silico screening permits the selection of high-affinity malaria antibodies.

## Introduction

The binding of an antibody to its cognate antigen is central to the application of antibodies as mediators of the humoral immune response and as biotherapeutic and prophylactic agents. *In vivo* a tightly regulated process of micro-anatomic evolution within the germinal center permits affinity maturation—driving low affinity precursors to acquire increasingly higher affinity for antigen through mutation and selection. In the case of antibodies targeting neutralizing epitopes, high-affinity variants have been shown to increase neutralization potential secondary to gains in affinity^1–3^.

In contrast, recent work has shown the potential of *in silico* approaches to the design of antibodies resulting in restoration of binding of clinically relevant antibodies to SARS-CoV-2^4,5^. A critical aspect of this emerging line of work is achieving “zero-shot” settings, in which computationally derived mutations are selected through *in silico* optimization alone. Despite the obvious utility of such approaches, it is well accepted that incorporation of increasing numbers of mutations is generally deleterious to the protein’s function, with only limited mutational paths to increased protein fitness^6–8^. Indeed, such zero-shot approaches have successfully introduced between four to seven mutations to increase antibody fitness.

Thus, a fundamental question remains unanswered—given a library of evolutionarily selected mutations, is it possible *in silico* to derive combinations of mutations associated with high protein fitness? Answering this question gives insight to the mutational “depth” that can be achieved during attempts to increase protein fitness. As a salient example, broadly neutralizing antibodies to HIV are known to possess both high mutational loads while retaining high fitness, as defined by viral neutralization^9–12^.

Here, we address this gap using two humanized mouse models that express BCRs specific to the circumsporozoite protein of *Plasmodium falciparum*, the most virulent etiological agent of malaria^13^. Globally, in 2022 estimated infections of malaria were 249 million with 608,000 deaths^14^. An emerging strategy for malaria prevention has been passive transfer of prophylactic antibodies, which have been shown to dramatically reduce parasitemia^15–17^. Although promising, these approaches have been marred by difficulties in achieving antibodies with high enough potencies to scale production in line with WHO guidelines^18^. Therefore, these mAbs present a critical case for extremely high affinity variants.

To determine affinity *in silico*, we hypothesized that information regarding overarching bond energies between antibody and antigen could be captured using structure prediction tools. By correlating antibody variant affinities to biochemical mechanisms of protein-protein binding, we demonstrate that relative affinities could be determined from predicted antibody-antigen structures. We then leveraged this strategy, developing a model inspired from the principles of the germinal center to explore sequence space as large as 10^47^ variants within hours. This model was then used to generate improved malaria prophylactic antibodies matured from each prototypic mAb with mutational loads of up to 31 amino acid substitutions demonstrating substantial ability to add mutations and improve antibody affinity.

## Results

### Number of protein-protein interactions in predicted CIS43 antibody:antigen complexes captures relative affinity rankings

CIS43 is a potent anti-CSP antibody isolated from a human volunteer immunized with an attenuated whole sporozoite vaccine and has been shown to be effective the prevention of malaria in endemic regions^15,19^. To improve the potency of this antibody, Kratochvil et al^20^. generated a transgenic mouse model that expressed the inferred germline of CIS43 (iGL-CIS43). Through immunization of this mouse line coupled with bioinformatic sieving of somatically mutated variants, the recombinant D3 variant was isolated. All three of these variants (iGL-CIS43, CIS43, and D3) primarily target the junctional epitope of *Plasmodium falciparum* CSP (PfCSP) with varying degrees of affinity^20^ (**Fig. 1A**). Obtaining variants with increased affinity is important to decrease the dose necessary for prophylaxis, which will ultimately increase the chance for deployment in resource-limited settings. Thus, we sought to understand whether structure prediction tools could offer insight into antibody:antigen affinity.

**Figure 1:**
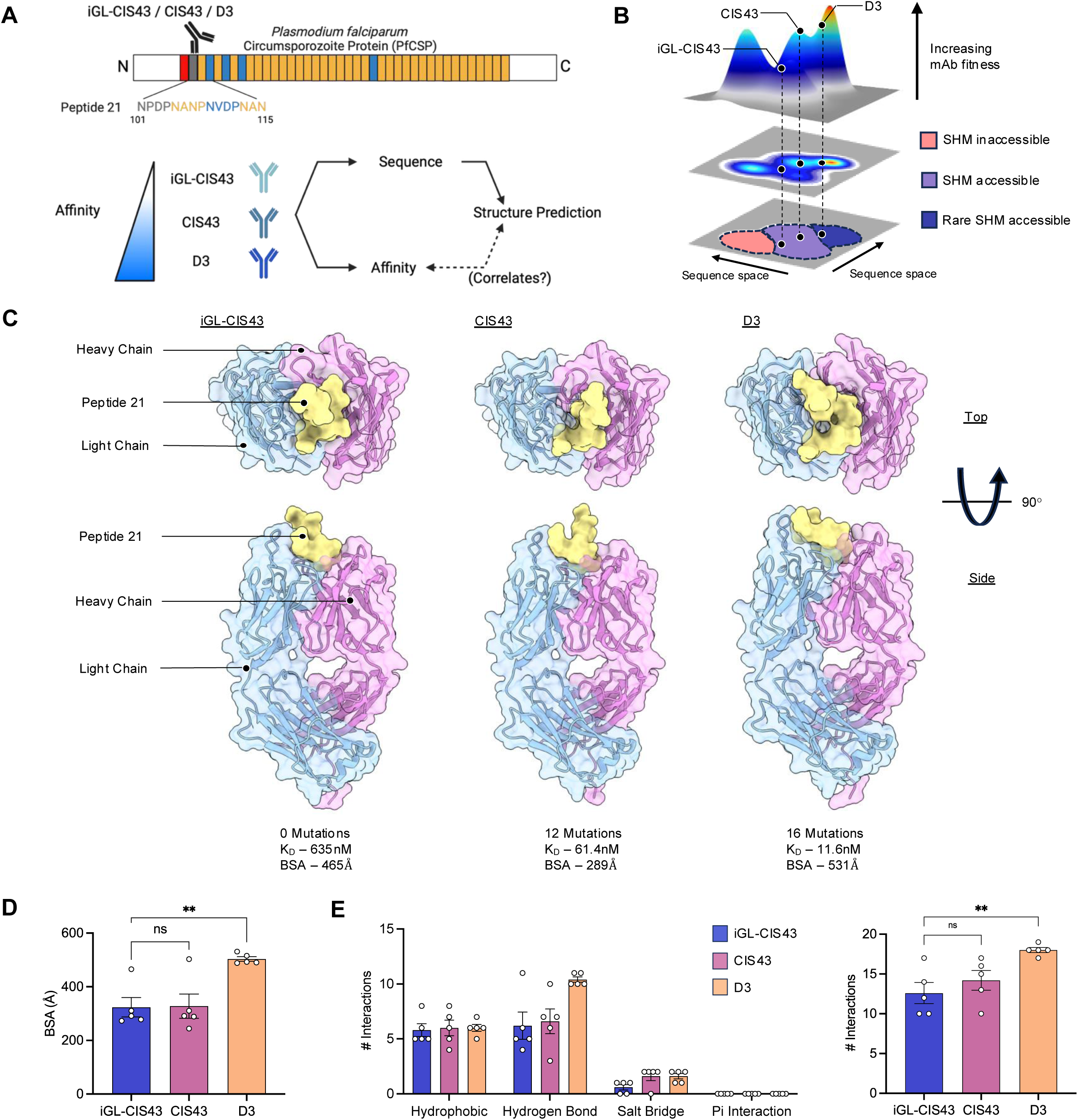
Fitness of CIS43 variant mAbs is captured by increasing number of intermolecular interactions between antibody:antigen. **(A)** Schematic demonstrating binding of CIS43-variant mAbs bind to PfCSP and the antigen peptide 21. iGL-CIS43, CIS43, and D3 variants were chosen to determine if affinity could be correlated to features of predicted structures. Reported Kd values are from Kratochvil et al. **(B)** Schematic depicting the hypothetical fitness landscape of CIS43 variant mAbs showing regions accessible to the process of somatic hypermutation (SHM). Regions not accessible to SHM are reflective of techniques such as random mutagenesis in yeast display libraries that have been shown to produce high fitness variants. Rare regions accessible to SHM are reflective of synthetic combinations of evolutionarily selected mutations in the germinal center. **(C)** Predicted structures of iGL-CIS43, CIS43, and D3 in complex with peptide 21. Shown structures are top ranked models from ColabFold analysis (see **Methods**). Experimentally determined K_D_ values are shown along with buried surface area (BSA) values determined from the predicted structures. **(D)** BSA values of peptide 21 derived from all 5 predicted models. **(E)** Shown are the number of interaction by type (left) and totals (right).

Through understanding of the relationship between affinity and sequence variants, we hypothesized that we could isolate more high affinity CIS43 variants. Notably, the potent variant D3 was not observed directly during immunization studies of Kratochvil et al., but rather, was comprised of a combination of somatic mutations isolated from those studies. Conceptually, we consider antibodies such as D3 to be rare variants that are somatic-hypermutation (SHM) accessible and hypothesize that other such variants with increased fitness exist but are difficult to elicit in a single sequence by SHM alone (**Fig. 1B**).

To determine whether structure prediction tools could inform isolation of these variants, we predicted the structures of iGL-CIS43, CIS43, and D3 in complex with the junctional epitope, peptide 21 (**Fig. 1C**) with ColabFold using AlphaFold-multimer^21,22^. Using all five outputs per antibody:antigen pair, we calculated the buried surface area (BSA) of peptide 21 in the antibody:antigen complex. Surprisingly, although we found that the high affinity variant (D3) possessed the greatest BSA, there was no difference between the low affinity (iGL-CIS43) and moderate affinity (CIS43) (**Fig. 1D**).

We next interrogated the type and number of protein-protein interactions between antibody and antigen. When stratified by type, we found that D3 contained the most hydrogen bonds, while both CIS43 and D3 contained equivalent number of salt bridges, and all three variants had equal number of hydrophobic interactions (**Fig. 1E, left**). Interestingly, when these interactions are summed, there appeared to be a stepwise increase in total interactions from low, medium, to high affinity (**Fig. 1E, right**). This suggested that the number of predicted interactions could covary with CIS43 variant affinity to peptide 21.

### Contact score correlates significantly with affinities of 38 iGL-CIS43 variants and can be used to construct novel high affinity variants

To determine whether the number of interactions indeed correlated with affinity, we sought to increase the number of antibody:antigen pairs. To do so, we analyzed 35 additional CIS43 variants that were identified during immunization studies^20^ (**Fig. 2A**). Affinities for these antibodies span two orders of magnitude and thus could be used to interpolate between iGL-CIS43, CIS43, and D3. For each of these antibodies we computed the predicted structure of the antibody:antigen complex and analyzed whether BSA, other computational metrics (predicted template modelling [pTM], interface predicted template modelling [ipTM], predicted aligned error [PAE], and predicted local distance difference test [pLDDT]) which measure global and local confidence in the predictions, or the overall number of interactions correlated with the corresponding affinities of the CIS43 variants. Surprisingly, we found that the number of interactions alone was sufficient to explain 49% of the variance within the dataset, which was more than double the next highest rank, pLDDT at 23%. (**Fig. 2B**). We then reasoned that there are differences in the relative strength of each type of interaction and thus weighting of the respective interactions would improve the correlation. Indeed, the derived metric “Contact Score,” was able to account for 63% of the variance in the dataset (**Fig. 2B**). Derivation of this metric can be found in **Methods**.

**Figure 2:**
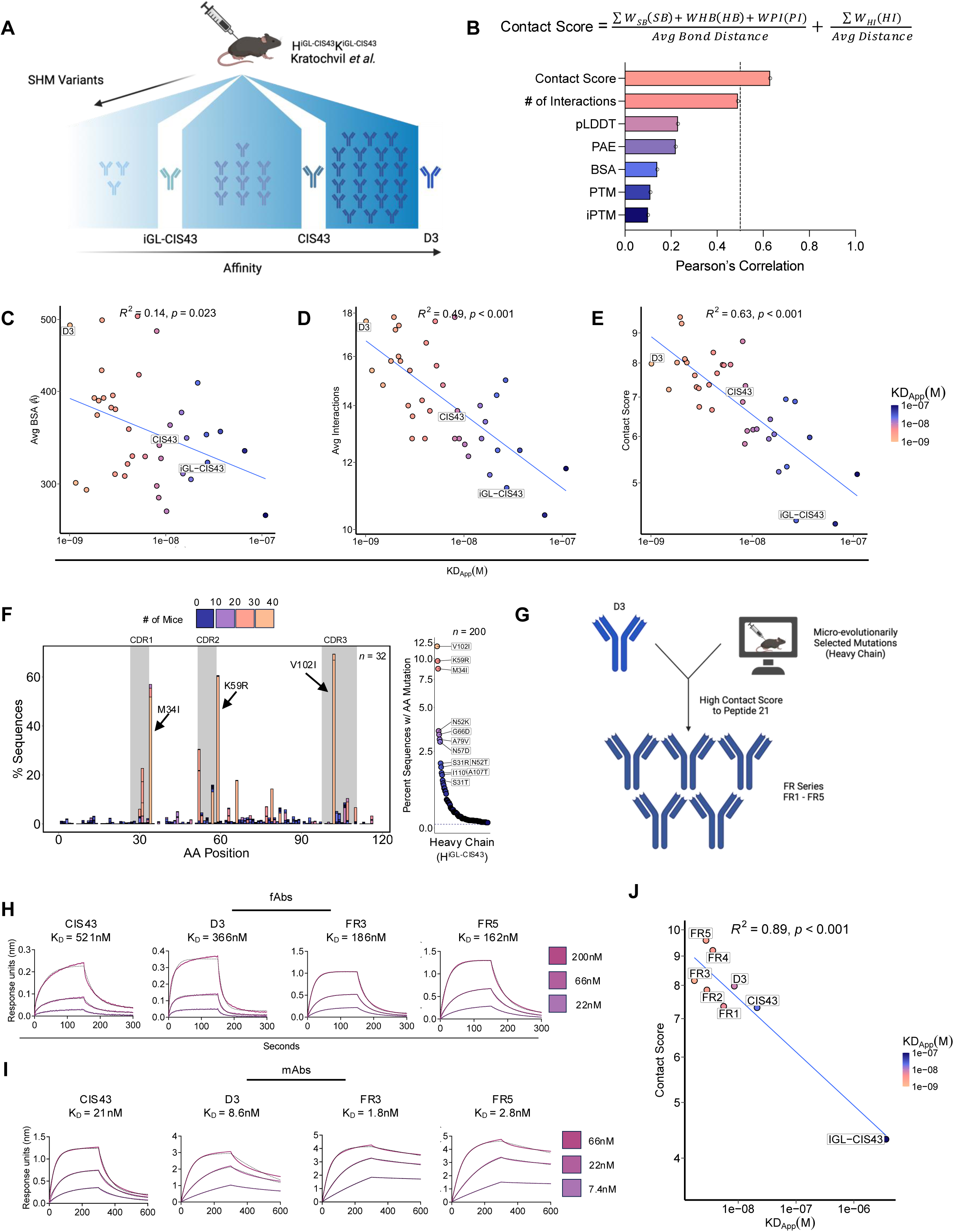
Contact Score is correlated to affinity of CIS43 variants and can be used to create novel high affinity variants. **(A)** Schematic depicting origin of 35 somatically mutated CIS43 variants described in Kratochvil et al. the affinity of which interpolates between iGL-CIS43, CIS43, and D3. **(B)** Pearson’s correlation for various metrics from predicted structures of 38 CIS43 mAbs to respective affinities to peptide 21. Contact score is defined by weighted sums of different types of intermolecular interactions divided by their corresponding average bond length (see **Methods**). SB – salt bridge, HB – hydrogen bond, PI – pi-interaction, HI – hydrophobic interaction. **(C–E)** Shown are the relationship between the average BSA **(C)**, average number of interactions **(D)**, and average Contact Score **(E)** to affinities of the 38 CIS43 variants to peptide 21. R^2^ and p-values are shown above. **(F)** Amino acid mutation frequencies of iGL-CIS43 sequences following immunization with peptide 21. Data are reflective of 38 independent mice and are colored to identify amino acid mutations shared across mice. Plot at right depicts frequency of amino acid mutations following removal of duplicate (clonal) sequences. **(G)** Schematic depicting generation of FR-series variants. Mutations identified in **(F)** were added to the heavy chain of the D3 variant and were selected for high Contact Scores. **(H–I)** Affinity measurements by biolayer interferometry of select FR-variants in both bivalent monovalent (Fab) **(H)** and full monoclonal antibody (mAb) **(I)** forms. **(J)** Correlation of FR-series variants to affinities as in **(C–E)**.

Examining the correlation between affinity and BSA, average number of interactions, and contact score reveals that while BSA can capture information about the affinity, there was little predictive value for both high and low BSAs (R^2^ = 0.14, p < 0.05) (**Fig. 2C**). In contrast, the average number of interactions demonstrated a clear stepwise relationship, with higher number of interactions corresponding to higher affinity antibodies (R^2^ = 0.49, p < 0.001) (**Fig. 2D**). Further improvements in the spread of the correlation were obtained by use of the Contact Score, which saw affinities closely approximating the regression line, although outliers were still present (R^2^ = 0.63, p = 3.4 x10^-9^) (**Fig. 2E**).

Next, we considered using the Contact Score to derive novel variants. To do so efficiently, we first sought to identify micro-evolutionarily selected mutations found from 11 to 95 days following first antigen exposure during immunization experiments in mice with B cells expressing transgenic iGL-CIS43 (**Fig. S1**). We reasoned that mutations that were selected across individual mice were likely key mutations that confer an affinity advantage while minimizing deleterious fitness effects. From 1,134 iGL-CIS43 heavy chain sequences isolated from 32 independent mice immunized with immunogens containing the binding epitope of peptide 21, we tabulated all amino acid substitutions found across the heavy chain VDJ regions. Unsurprisingly, we found concerted selection for key mutations V102I, K59R, and M34I (**Fig. 2F**). These mutations were highly enriched within the sequence set, accounting for >40% of all sequences and occurring in nearly every mouse. Both V102I and M34I mutations occurred in tandem with an activation-induced deaminase (AID)-associated hotspots, while K59R was not, suggesting selection for both frequent and infrequent mutations. We then considered that clonal effects within mice could result in confounding effects, thus we removed all duplicate sequences from the sequence set and calculated the frequency of each mutation. We found enrichment of the same mutations, with other key mutations emerging from background (**Fig. 2F**).

Using D3 as a template, we created five variants that expressed additional micro-evolutionarily selected mutations and still retained a high Contact Score to peptide 21 (**Fig. 2G**). Notably, some variants also incorporated a beneficial mutation (P104K) from an improved CIS43 variant, P3-43^23^. These variants, collectively termed the FR-series (**Fig. S2**), were expressed and their affinities derived to peptide 21. We first expressed a subset of FR variants as monovalent single-domain fragment antigen-binding (Fab) regions and assayed them to confirm that binding improvements were due to genuine affinity changes rather than avidity effects. As Fabs, FR3 and FR5 displayed KD values of 186 nM and 162 nM, respectively, compared to 366 nM for D3 and 521 nM for CIS43 (**Fig. 2H**), representing ∼3-fold improvement over CIS43 and ∼2-fold over D3. We then measured the set as bivalent full-length antibodies and found that FR3 and FR5 mAbs also showed increased affinity to peptide 21, with an approximately 4-fold improvement relative to D3 (**Fig. 2I**).

We next sought to confirm that the relationship between the affinity of the novel FR series variants aligned with the relationship observed prior between affinity and Contact Score. Indeed, we observed a strong correlation between these variants, D3, CIS43, and iGL-CIS43 (R^2^ = 0.89, p < 0.001) (**Fig 2J**). These results suggest that novel variants can be efficiently identified from combinations of selected mutations when screened by the Contact Score metric.

### The iGC algorithm for iGL-CIS43 navigates sequence space and independently produces high-affinity mAbs

Having identified variants with increased affinity relying on contact score as an *in silico* screen, we next sought to generate an automated pipeline for exploring the vast space of possible CIS43 variants, including mutation data of the light chain as well (**Fig. S3**). We identified a pool of 200 heavy chain mutations and 113 light chain mutations from iGL-CIS43 expressing mice for a combined combinatorial space of >3 x 10^47^. To efficiently explore this sequence space, we drew inspiration from the biology of the germinal center, in which cells compete for acquisition of antigen by mutating and selecting for the highest affinity variants among their peers. Similarly, using Contact Score as a proxy, we developed the in silico germinal center (isGC) pipeline, which follows the same principles as the biologic germinal center but with Contact Score as the metric by which sequences are selected (**Fig. 3A**). The isGC pipeline is composed of three branches: mutation, prediction, and selection. Each iteration through these processes is considered a cycle. The mutation step is performed using a weighted matrix of observed *in vivo* mutations, with those mutations occurring most frequently selected the most often according to the mutation plot shown in **Fig. 2F**. By doing so, we hypothesized that the enrichment of beneficial mutations would lead to less deleterious outcomes, in a strategy similar to work by Patsch et al. for enzyme design^8^. These mutated sequences are then passed into the prediction step, in which a predicted structure is generated for the sequence complexed with antigen. In the selection step, those sequences associated with the highest contact score are passed into the next cycle of mutation and selection.

**Figure 3:**
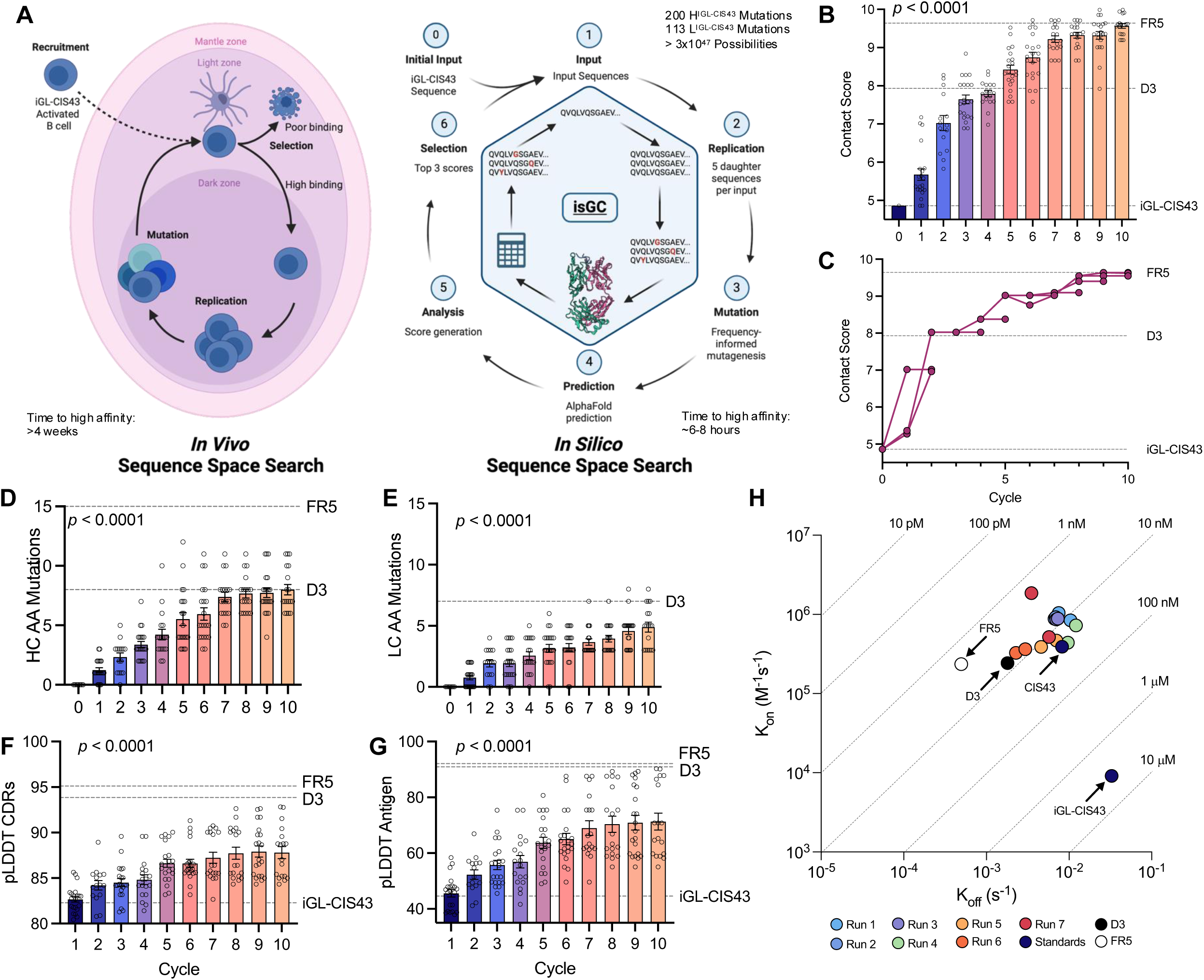
The isGC algorithm for iGL-CIS43 navigates sequence space and independently produces high-affinity mAbs. **(A)** Schematic of the biological mechanism by which B cells survey sequence space in the germinal center (left) as compared to the isGC pipeline (right). Created using BioRender. **(B)** Bar plot depicting the change in Contact Score as sequences progress through multiple cycles. Shown are the top three scores at each cycle for seven independent runs. **(C)** Connected scatter plot depicting the relationship between the top three Contact Scores per cycle. Shown in color is a single individual run through the isGC pipeline. Six other runs are shown in grey scale to demonstrate the distribution of scores over subsequent cycles. **(D–E)** Bar plots depicting the number of mutations gained in the heavy (**D**) and light (**E**) chains following progression through the isGC pipeline for 10 cycles. **(F–G)** Bar plots depicting average pLDDT values for antigen (**F**) and antibody CDRs (**G**) following progression through the isGC pipeline for 10 cycles. **(H)** BLI measures of composite K_on_ and K_off_ values for selected isGC-derived monoclonal antibodies to peptide 21. Two representative sequences from cycle 10 of each run of the isGC are presented. Isoaffinity lines are drawn for reference.

Starting from the iGL-CIS43 sequence, we performed the pipeline seven times with a limit of 10 cycles of mutation and selection (**Fig. S3**). As expected, sequences exhibited improved Contact Scores over each of these cycles (**Fig. 3B–C**). Notably, each of the runs of the isGC appeared to reach an asymptote of a contact score of ∼10, suggesting a physical limit of affinity to antigen^24^. Indeed, the observed contact scores exceeded D3 but were like FR5. Analysis of the number of heavy and light chain mutations per cycle yielded similar results, with the number of mutations steadily increasing with each cycle before plateauing (**Fig. 3D–E**). Notably, while these sequences reached mutation levels similar to D3, they did not reach that of FR5. We next examined the effect of these mutations on the quality of predicted structures. Unexpectedly, we found that with increasing Contact Scores, generally disordered regions, including CDR loops and antigen, significantly increased in AlphaFold’s per-residue confidence metric, pLDDT (**Fig. 3F–G**) (*p* <0.001). This suggested that the mutations added during the isGC, both select for increased Contact Score as well as increased AlphaFold’s confidence in regions considered to be difficult to model^25,26^. This was also mirrored in the pLDDT values of, in order of increasing affinity, iGL-CIS43, CIS43, D3, and FR5 (**Fig. S4**).

From each of these runs, we selected two sequences with the top Contact Scores, expressed them, and performed BLI apparent affinity measures to peptide 21. As suggested by the contact scores, we found that all antibodies expressed possessed affinities in the range of D3, with half possessing affinities greater than D3, and in one case we identified a variant with affinity similar to that of FR5 (**Fig. 3H**). Affinities of these variants ranged from 22 nM to 1.8 nM, indicating a 150-to-2000-fold improvement over the starting sequence, iGL-CIS43. Further analysis of kinetics demonstrated that each variant selected for overall affinity, rather than biasing uniformly towards K_on_ or K_off_ (**Fig. 3H**). This is expected given the maturation of iGL-CIS43 to the mature CIS43 followed a similar trajectory. Despite this, we did not observe variants that had achieved off-rates comparable to FR5 or D3, suggesting that by starting at iGL-CIS43, the gains in affinity may be initially driven by gains in association rates as compared to maturation of a more constrained, high affinity variant, such as D3.

Separately, we asked whether the selection of Contact Score during each cycle was meaningfully selecting high affinity variants. To answer this question, we generated eight pairs of antibodies originating from each of the seven runs of the isGC. These pairs were matched so that a high Contact Score variant selected at the end of a cycle (parent sequence) was matched with a daughter sequence from the next cycle which had additional mutations added and was determined to have a lower Contact Score than the originating parent sequence (**Fig. S5**). We expressed and tested these 16 variants and found that for 7 of 8 pairs, the decrease in contact score correctly predicted the decrease in affinity as assayed by serial dilution ELISA (**Fig. S6**). Importantly, we found this effect occurred in the presence of 1–2 additional mutations, demonstrating that the Contact Score correctly identified when non-optimal, yet micro-evolutionarily selected, mutations were added to a given variant backbone.

### The isGC pipeline using FR5 produces variants with greater affinity not predicted by AlphaFold pLDDT score

Given the disparate results between the FR-series and the isGC output, we hypothesized that it could be possible to further enhance the affinity of FR5 by using it as input into the isGC algorithm. Specifically, we noted that the isGC variants were selected for high K_on_ and high K_off_ while the FR5 variant had decreased K_off_. We suspected that by starting at FR5, we may be able to constrain the evolution of the antibody:antigen binding to decreased K_off_. We also questioned whether total mutation load could provide an alternative explanation for the differences observed between FR5 and the isGC variants. Thus, we performed three runs of the isGC using FR5 as starting input. From these runs, we isolated nine variants, which we term collectively as the SFR series (**Fig. 4A, Fig. S2**). Within this set, four variants were selected for high Contact Scores (> FR5), one for high Contact Score and high mutation load, and four who had high mutation loads but retained a moderate Contact Score (similar to D3).

**Figure 4:**
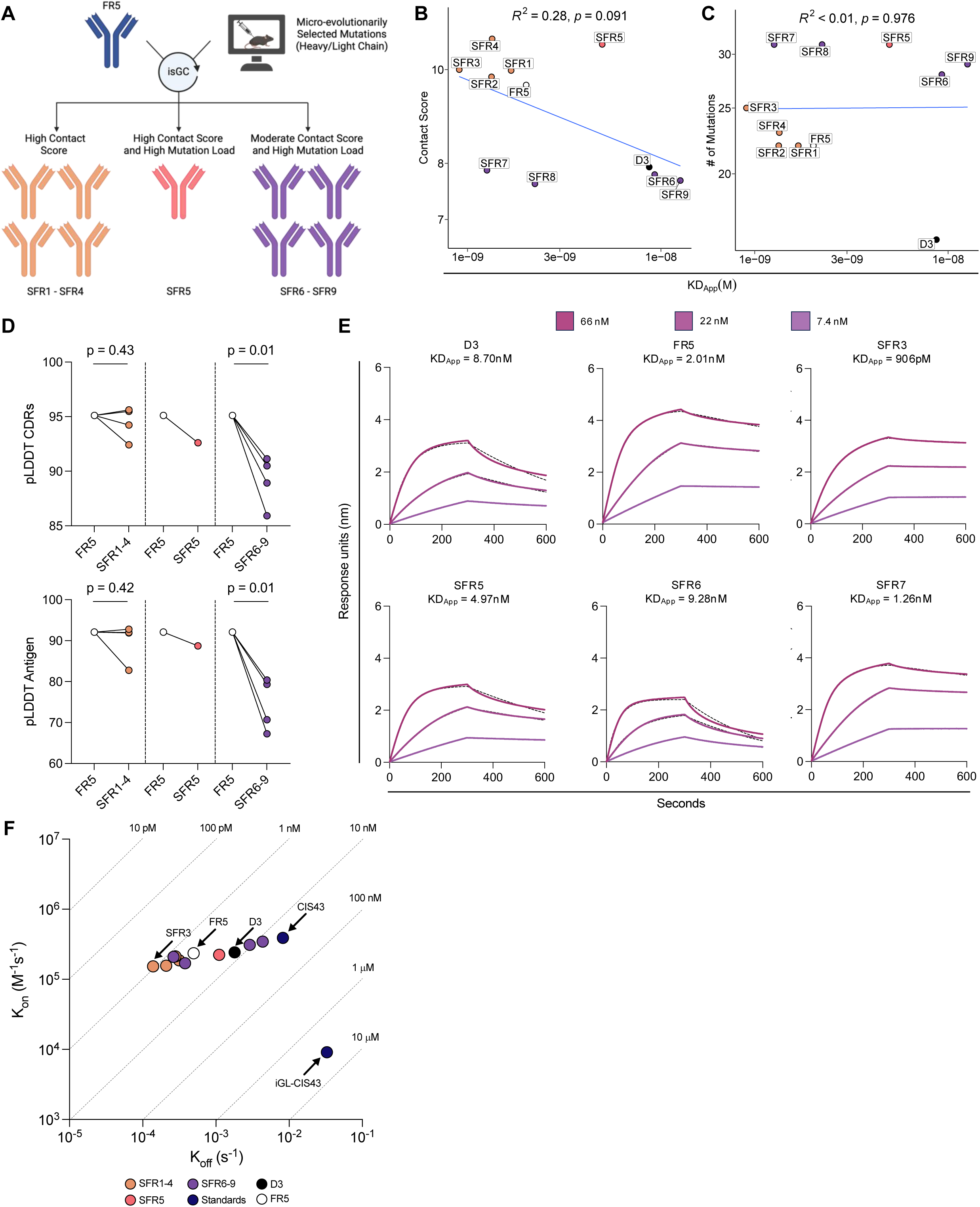
The isGC pipeline using FR5 produces variants with greater affinity not predicted by pLDDT score. **(A**) Schematic of the generation of SFR-variants. Using FR5 as input into the isGC, three classes of variants were selected for expression: those with high contact scores, those with high contact scores and mutation loads, and those with high mutation loads but moderate contact scores. **(B– C**) Correlation of affinity to peptide 21 and contact score (**B**) or mutation load (**C**). Data are depicted as in **Fig 2C**. **(D)** Relationship between pLDDT of CDRs (Top) and antigen (Bottom) of SFR variants compared to the parental FR5 variant. **(E)** Affinity measurements by BLI to peptide 21 of SFR variant mAbs. Data are presented as in **Fig. 2I**. **(F)** Scatter plot depicting the composite K_on_ and K_off_ values for SFR variants. Isoaffinity lines are drawn for reference and are depicted as in **Fig. 3H**.

As before, we expressed the SFR variants and measured apparent affinities to peptide 21 via BLI. We again compared affinities to Contact Scores and found a correlation that approached significance (**Fig. 4B**) (R^2^ = 0.28, p = 0.091). We found a series of improved SFR variants within the subset selected solely on high Contact Scores, with our best variant being SFR3 which exhibits 10-fold increase in affinity over D3. Interestingly, we found that SFR variants SFR5, SFR7, and SFR8 were outliers. SFR5 was selected for having both high mutation loads and a high Contact Score, suggesting that the Contact Score can be errant at extremely high mutation loads. Similarly, SFR7 and SFR8 represent the inverse, with moderate contact scores but very high affinity despite high mutation loads. Correlation of mutation loads to affinity confirmed that mutation load alone does not covary with affinity (**Fig. 4C**).

We next looked at AlphaFold’s confidence metric pLDDT for both antigen and for CDRs to understand whether Contact Score continued to covary with confidence metrics for predicted structures. We found that for both antigen and CDRs, SFR variants with high Contact Scores did not have pLDDT values significantly change after selection in the isGC (**Fig. 4D**) (p > 0.05). Conversely, those that have moderate Contact Scores all had significant decreases in confidence metrics (p = 0.01). This result was intriguing as it suggests, in tandem with the affinity results, that there are “holes” in the predictive capacity of protein prediction models, especially at very high mutation loads, and could inform future refinement of such models^27^.

### The isGC pipeline can accommodate and mature complex antibody-antigen and antibody-antibody interactions simultaneously

We next sought to extend application of the isGC algorithm to improve the affinity of L9, another potent anti-malarial antibody undergoing clinical trial^17^. Like CIS43, L9 targets CSP present on the circumsporozoite stage of the parasite. Unlike CIS43, L9 primarily targets the minor repeat region, which is composed of three NPNV repeats (**Fig. 5A**). To run the isGC, we identified micro-evolutionarily selected mutations in a data set from an upcoming manuscript^28^ wherein humanized knock-in mice bearing the mature L9 heavy and light chain were immunized with the immunofocusing antigen peptide 22, which contains two NPNV repeats.

**Figure 5:**
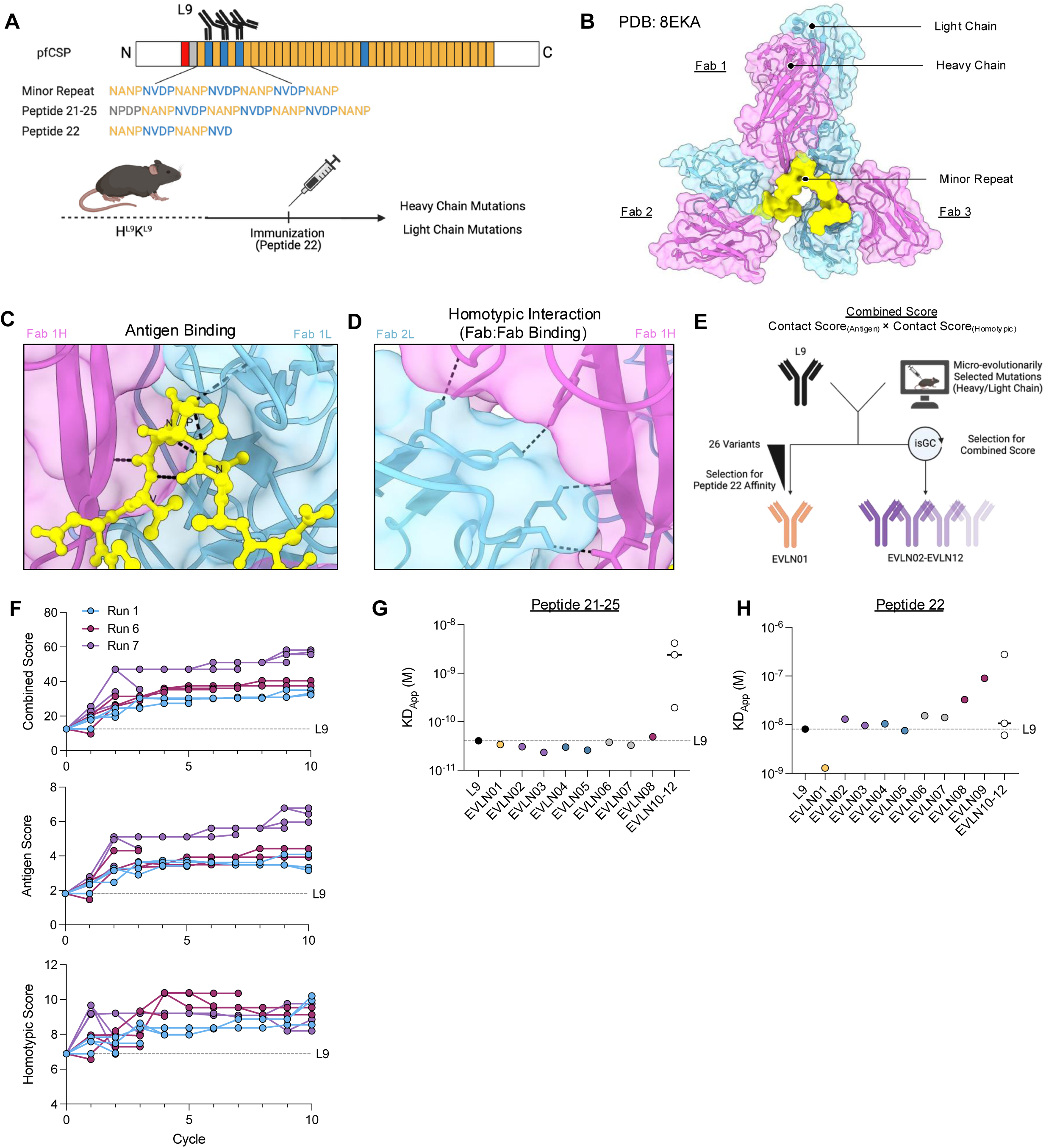
The isGC algorithm applied L9 demonstrates the capacity to accommodate complex antibody:antigen interactions. **(A)** Schematic of the anti-CSP antibody L9. Shown is the schema for the H^L9^K^L9^ humanized mice and immunization with cognate antigen peptide 22. The relationship between peptide 22 and PfCSP is shown. Additionally, as L9 is known to form a larger quaternary structure on PfCSP, the *in vivo* target (minor repeat) and its corresponding peptide fragment (peptide 21–25) is also shown. The binding epitope of L9 is shown for each of these as the underlined amino acids. **(B)** Visualization of the L9:Ag complex. Shown are three Fab arms that each bind the NPNV motif on the minor repeat. **(C–D)** Shown are the major intermolecular interactions present in the L9 complex including L9:antigen interactions **(C)** and homotypic L9 interactions **(D)** in which there are contacts between adjacent Fabs. **(E)** Schematic describing the generation of the EVLN-series mAbs. EVLN01 was made by expressing L9 variants and selecting based upon the highest affinity to peptide 22. EVLN02–EVLN12 were made through the isGC with selection criteria for high Combined Score. This score is comprised of the Contact Score to antigen multiplied by the Contact Score of homotypic interactions. **(F)** Connected scatter plot depicting the Combined Contact Score (Top), the Antigen Contact Score (Middle) and the Homotypic Contact Score (Bottom) over seven L9 isGC runs of 10 cycles. Selected runs are highlighted. **(G–H)** Apparent affinities for selected isGC-derived L9 sequences to peptide 21-25 (**G**) and peptide 22 (**H**). Colors correspond to runs highlighted in (**F**).

The structure of L9 is unique with a complex quaternary structure (**Fig. 5B**). The complex is comprised of typical antibody:antigen binding (**Fig. 5C**) as well as homotypic interactions (**Fig. 5D**) in which individual Fabs form contacts with neighboring Fabs. Previous studies have shown that the neutralizing potential of L9 is derived from these two key intermolecular interactions^29–31^. Thus, maturation of this antibody would require optimization of both antibody:antigen interactions as well as antibody:antibody interactions.

To apply the isGC pipeline to this structure, we derived a modified Contact Score: in addition to deriving the Contact Score to antigen, we determined the Contact Score between the homotypic interface and multiplied these values to generate a Combined Score to enforce selection of both (**Fig. 5E**). The Combined Score was treated as before in the isGC pipeline, with those sequences associated with high scores proceeding to additional rounds of mutation and selection. Expressed variants were designated as the EVLN series and includes a single variant, EVLN01, that was optimized with micro-evolutionarily selected mutations, but without use of the isGC, for high affinity to peptide 22. As before, the Combined Score increased through successive cycles (**Fig. 5F**). Splitting the combined score into its respective components, we found that score to antigen also increased with each cycle, while the homotypic score increased initially but tended to vary in tandem with the increases seen in antigen score. As with CIS43 mAbs, we also observed increases in AlphaFold metrics of low confidence regions (CDRs and antigen) (**Fig. S6**).

We expressed these predicted variants and measured affinity to both peptide 21-25, which contains all three NVDP motifs that the predictions were modeled on, and to peptide 22. We found that binding to peptide 21-25 was improved in three of the seven runs, maintained in one, and lost in another three as compared to L9 (**Fig. 5G**). Given that these mutations were derived in the context of peptide 22 immunization, we also assayed against peptide 22 for apparent affinity. We found that most sequences lost affinity to peptide 22, with a single exception (**Fig. 5H**). The generation of the higher affinity variants from an alternative malaria antibody demonstrated that the isGC pipeline could efficiently incorporate complex antibody-antigen and antibody-antibody interactions.

## Discussion

In this study, we utilized a library of micro-evolutionarily selected mutations and demonstrated that such a library could be coupled with protein structure prediction to design high affinity. We showed that affinity is tightly linked with a metric derived from intermolecular interactions and can be used to drive affinity selection both in a simple 1:1 antibody-antigen model (CIS43) as well as in a complex 3:1 antibody-antigen model (L9). We then used this metric to design a computational pipeline that could select for variants with increased affinity from a vast computational space within hours. Altogether, this study demonstrates a proof of principle that efficient in silico maturation of antibody variants is possible and that such maturation can engage significant mutational depths without compromising antibody affinity.

In the time since the development of the AlphaFold architecture, there has been a surge in protein prediction tools, with increasing ability to predict the native structure of a protein or protein complex^32^. In tandem, significant work has attempted to computationally determine protein binding affinity^33,34^. However, despite progress in both arenas, antibody:antigen complexes have proven difficult to model structurally, and difficult to predict affinities for^33–35^. Due to the large therapeutic potential of antibodies as biomolecules, there is a pressing need to understand how antibodies engage with their targets and subsequently how tightly they bind. Thus, recent studies have paired protein prediction models alongside other computational approaches to derive affinity for antibody:antigen pairs^36–39^.

Although there have been studies that pursued this approach, the methods thus far have suffered from high computational expense^5^. Here, we demonstrate that by restricting to micro-evolutionarily selected mutations, we can leverage a less computationally expensive tool (AlphaFold-Multimer / ColabFold) enabling rapid interrogation of large swaths of mutational space while improving antibody fitness. This suggests that AlphaFold has the capacity for interrogating mutational space, even in the context of challenging targets such as L9. If true, understanding the determinants of this effect could drive further work for *in silico* optimization of antibody:antigen pairs as well as other classes of proteins writ large, given the ability of AlphaFold to model these to a high degree of confidence.

Of particular interest is the relationship between affinity and pLDDT as shown from this work. Given that the ability of AlphaFold to derive structures is linked with co-evolutionary signal generated from multiple sequence alignments (MSA)^21,40^, it would not be expected that introduction of point mutations would significantly change the MSA between low affinity/pLDDT variants. Despite this, we observe increases in pLDDT in tandem with Contact Score. It has been previously shown that intrinsically disordered regions (IDRs) generally result in a low pLDDT score but in instances where these regions become structured after binding to a target protein, the pLDDT increases^41,42^. In this work, it appears that through the introduction of mutations, these hard-to-model regions such as the CDRs become “locked” in a relatively constrained way such that the confidence (pLDDT score) in the prediction of these sites increase dramatically and suggests that local optimization of the pLDDT score can be leveraged for computational refinement and design of similar antibodies.

In the present study we identified multiple high affinity antibody variants of two potent anti-malarial antibodies in clinical trials, CIS43 and L9. While affinity does not always correlate with potency, analyzing the in vivo protective capacity of the variants generated in this study may help determine the extent to which they correlate in this system. Multiple techniques have been used to identify protective antibodies and antibody variants of this class including yeast display, knock-in mouse models, and screening from immunized human donors^19,20,43–45^. The higher affinity variants identified here suggest that protein prediction models can contribute to efforts to eradicate malaria and could be leveraged for the rapid affinity enhancement of other clinically relevant antibodies.

## Methods

### Structure prediction and scoring of predicted structures

Structural prediction of antibody:antigen pairs was performed using localColabfold^22^ with AlphaFold-Multimer_v3 weights^21^. All predictions were generated with the template and dropout flags enabled; all other options were left in their default settings. To model CIS43 variant antibody:antigen pairs, antibody variants were modeled as Fabs. For heavy chain sequence, inputs included the first domain of the heavy chain constant region of human IgG1. For light chains, inputs contained the constant region of the kappa light chain. Antigen for CIS43-variants were fixed as the PfCSP junctional peptide 21 (NPDPNANPNVDPNAN).

Derivation of the Contact Score was performed using a custom script written in R. In brief, for each antibody:antigen pair, localColabfold returned five predicted structures. For each, the predicted structure was interrogated for predicted polar and nonpolar interactions. Polar interactions were then stratified and weighted by type with increasingly polar side chain interactions weighed more heavily. Each weighted interaction was summed and normalized by the average predicted bond distance of all polar interactions. Similarly, nonpolar interactions were surveyed using PLIP^46^. Each of these interactions was similarly weighted and normalized by average predicted bond length of all non-polar interactions. A Contact Score is derived from each of the five predicted models and then averaged yielding the final Contact Score for the antibody:antigen pair.

For L9-variant modeling, antibody:antigen pairs consisted of a single antigen, peptide 21-25 (NPDPNANPNVDPNANPNVDPNANPNVDPNANP), along with three variable region-only heavy and light chain L9-variant sequences. To derive the “Combined Contact Score” metric for these structures, separate contact scores were generated for antibody:antigen interactions as well as those for antibody:antibody interactions. These values were multiplicatively combined.

### In Silico Germinal Center

The In Silico Germinal Center (isGC) pipeline utilizes a custom script written in R and Bash. The script is composed of three interdependent parts and accepts an input antibody:antigen sequence and a weighted substitution matrix of allowed mutations. For the work described here, substitution matrices were derived from the observed mutation frequences in knockin B cells following immunization experiments. Substitution matrix weights are derived from the frequency of observed mutations from the immunization data. In brief, input sequences are randomly selected to add up to 3 mutations in the antibody heavy chain and up to 2 mutations in the light chain. Mutations are selected from the provided substitution matrices. In total, five newly mutated sequences are generated and subsequently passed into localColabfold for structure prediction as described. Following prediction of these variants, each variant is scored as described above. The top three scoring sequences are retained and used as input in the next cycle of the isGC. This continues for the number of user-specified cycles.

### Affinity measurements by BLI

Biolayer interferometry (BLI) measurements of antibody:antigen binding affinity were performed on an Octet Red96e (fortéBio) utilizing streptavidin capture biosensors in black 96 -well plates (Greiner Bio One). Assays consisted of immobilization of biotinylated antigen for five minutes followed by a 60 second baseline in 1X Kinetics Buffer (Sartorius). Antigen was loaded in non-saturating conditions. Antigen-loaded biosensors were dipped into dilutions of full-length antibody or Fabs for five minutes and followed by a five-minute dissociation in 1X Kinetics Buffer. All assays were performed at 25°C with agitation. All Octet measures were adjusted to subtract baseline drift by measuring an antigen loaded sensor in 1X Kinetics Buffer. Binding traces were fitted globally with a 1:1 Langmuir model of binding. Data analysis was performed on Octet Software, version 9.0.

### Affinity measurements (ELISA)

High binding capacity 96-well plates (Nunc) were coated overnight at 4°C with 50ng/well of peptide 21 or peptide 22. Plates were washed three times with 0.05% Tween-20 in PBS (PBST), blocked with 3% BSA in PBS for one hour at RT and washed again three times with PBST. Antibodies were serially diluted three-fold and incubated for an hour at RT, washed three times, and incubated with Alkaline Phosphatase AffiniPure Goat Anti-Human IgG (Jackson ImmunoResearch) at 1:1,000 in PBS for one hour at RT. p-Nitrophenyl phosphate dissolved in water was used to detect signal. ELISA curves were generated in Graphpad Prism 10.

### Mouse lines and immunizations

Knockin mouse lines were previously published or are in publication by our group^20,28^. For all immunization experiments performed, immunogens were diluted in 100 μL PBS and mixed in equal volume of 2% alhydrogel for a total combined volume of 200 μL. All mice were immunized interperitoneally. All experiments were done with approval by the Institutional Animal Care and Use Committee (IACUC) of Harvard University and the Massachusetts General Hospital and conducted in accordance with the regulations of the American Association for the Accreditation of Laboratory Animal Care (AAALAC).

## Acknowledgements

We would like to thank all members of the Batista lab for their help and support, particularly Abigail Esposito, Usha Nair, and John E. Warner. We would also like to thank Azza Idris (Ragon), Peter D. Kwong (Columbia University), and Robert Seder (NIH), and their lab members for discussion and feedback on this manuscript. Thanks also to the Ragon Scientific Editing Platform for copyediting. This work was supported financially by National Institutes of Health NIAID R01 AI168114 and R01 AI151178 (to FDB); the Gates Foundation INV 009585 (to FDB), and flexible funding from the Ragon Institute (to FDB). This project has also been funded in whole or in part with Federal funds from the National Cancer Institute, National Institutes of Health, Task Order No. 75N91019F00135 under Contract No. 75N9101900024. The content of this publication does not necessarily reflect the views or policies of the Department of Health and Human Services, nor does mention of trade names, commercial products, or organizations imply endorsement by the U.S. Government.

## Author Contributions

GAD: conceptualization, data curation, formal analysis, investigation (affinity measurements), methodology (script generation; algorithm development), writing—original draft; BC: data curation, methodology (ColabFold pipeline establishment; script generation); JHK: data curation, investigation; MW: investigation; SK: data curation, investigation; FDB: conceptualization, funding acquisition, project administration, supervision, writing—review & editing

## Data Availability Section

All sequence data has either been or will be made publicly available in GenBank as part of the experimental source papers listed and can be provided to reviewers on request; this section will be replaced by GenBank reference numbers prior to publication. The underlying code may also be provided confidentially to reviewers on request and will be deposited prior to publication; this section will be replaced by a link and reference.

## Supplemental Figure Legends

**Figure S1:**
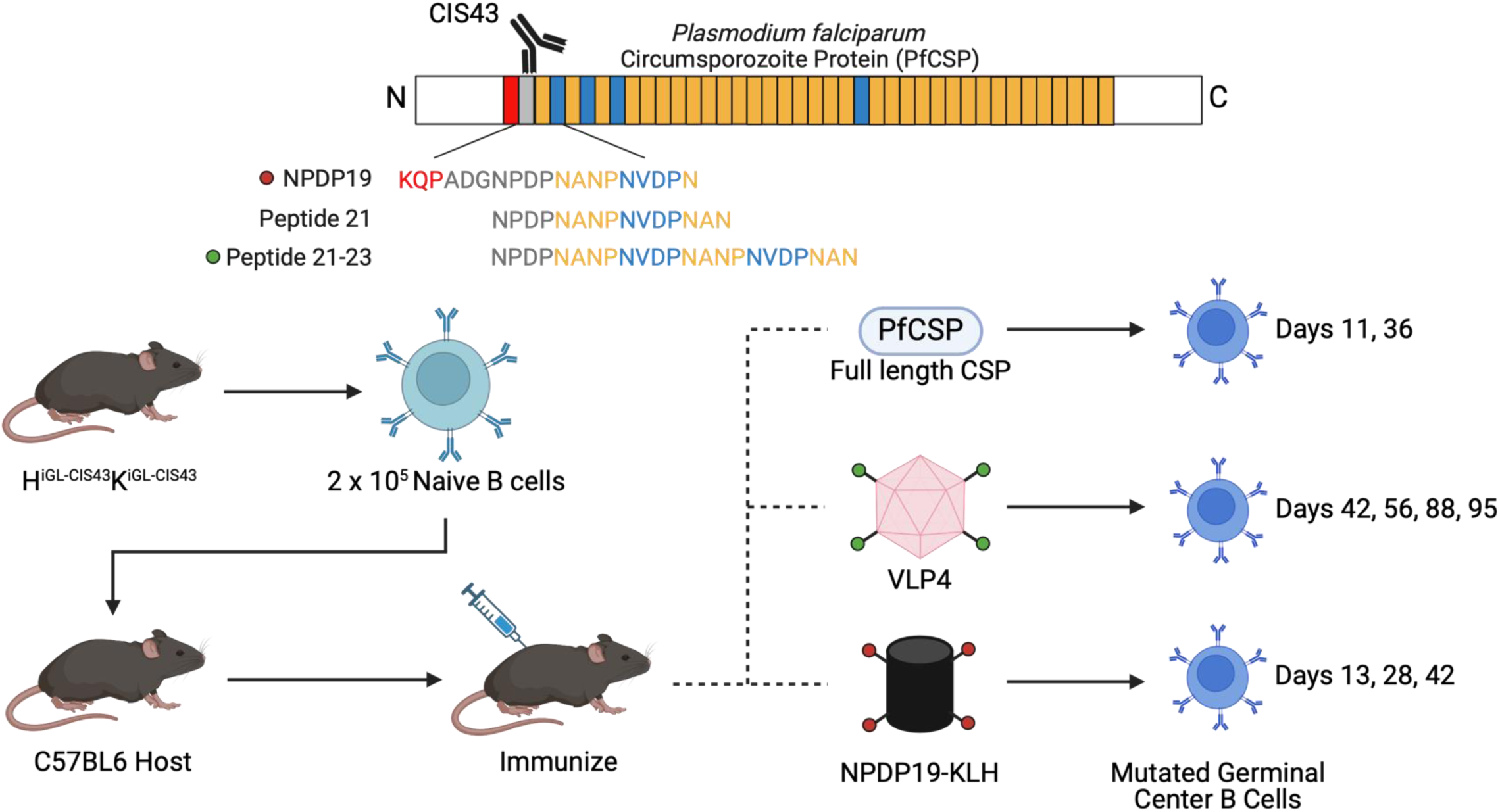
Overview of CIS43 experiments yielding mutation data used in isGC pipeline. Splenic B cells from H^iGL-CIS43^K^iGL-CIS43^ mice were harvested and 2 x 10^5^ B cells were transferred to C57BL6 host mice. These mice were immunized with either full length PfCSP, VLP4 (which encodes Peptide 21-23), or NPDP19-KLH. Cohorts of mice were sacrificed at the listed time points and germinal center B cells were collected by FACS for single cell sequencing.

**Figure S2:**
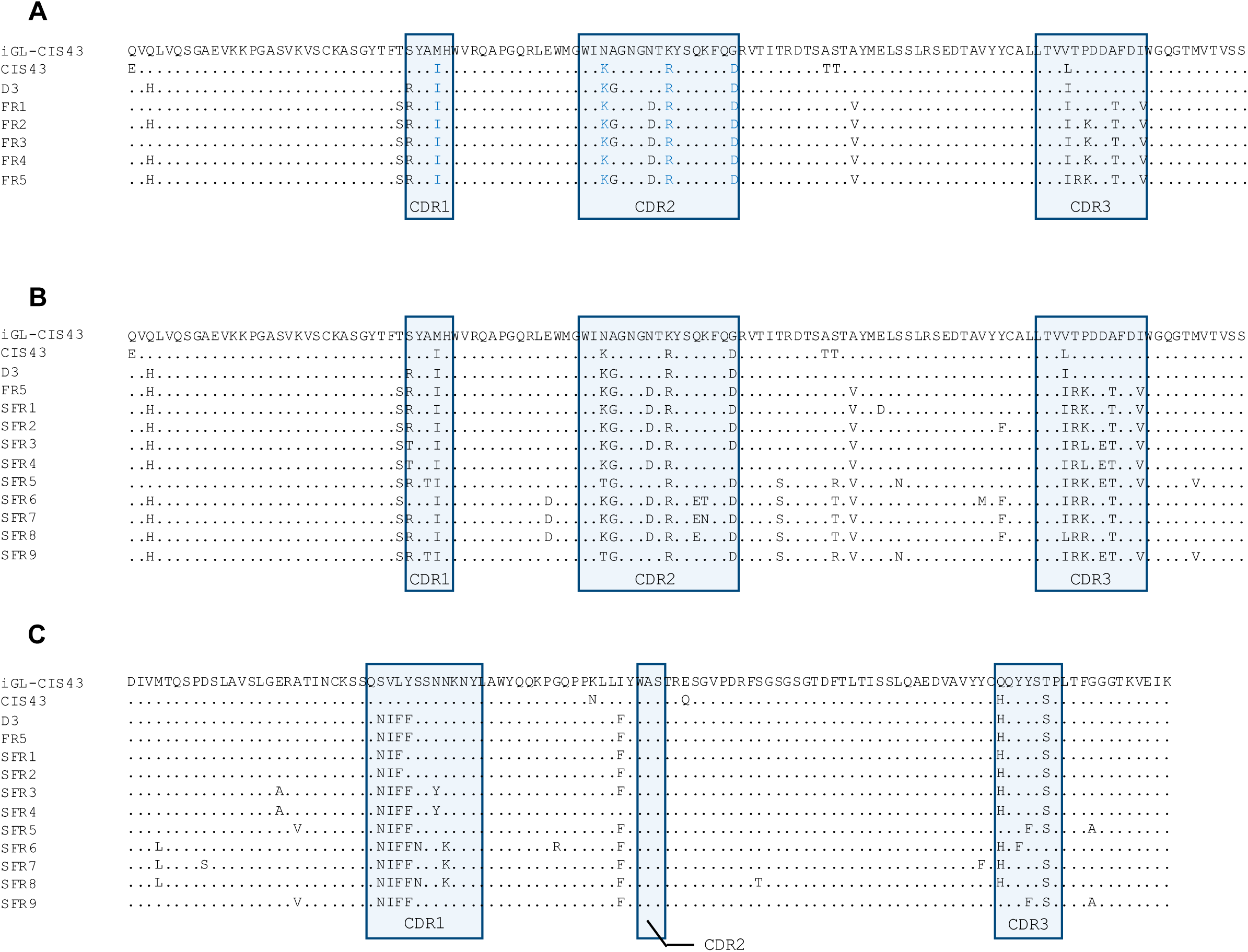
Sequences of FR and SFR series variants. **(A)** Shown are the heavy chain of iGL-CIS43, CIS43, D3, and the FR-series variants. Sequences are depicted with identities marked as dots with mutations shown as their respective amino acid. Mutations shared among all variants are shown in blue. Shaded regions demarcate the three heavy chain complementarity determining regions (CDRs). FR series variants share the same light chain as D3 (not shown). **(B–C)** Shown are the heavy **(B)** and light chain **(C)** sequences of IGL-CIS43, CIS43, D3, and the SFR-series variants. Data are presented as in **(A)**.

**Figure S3:**
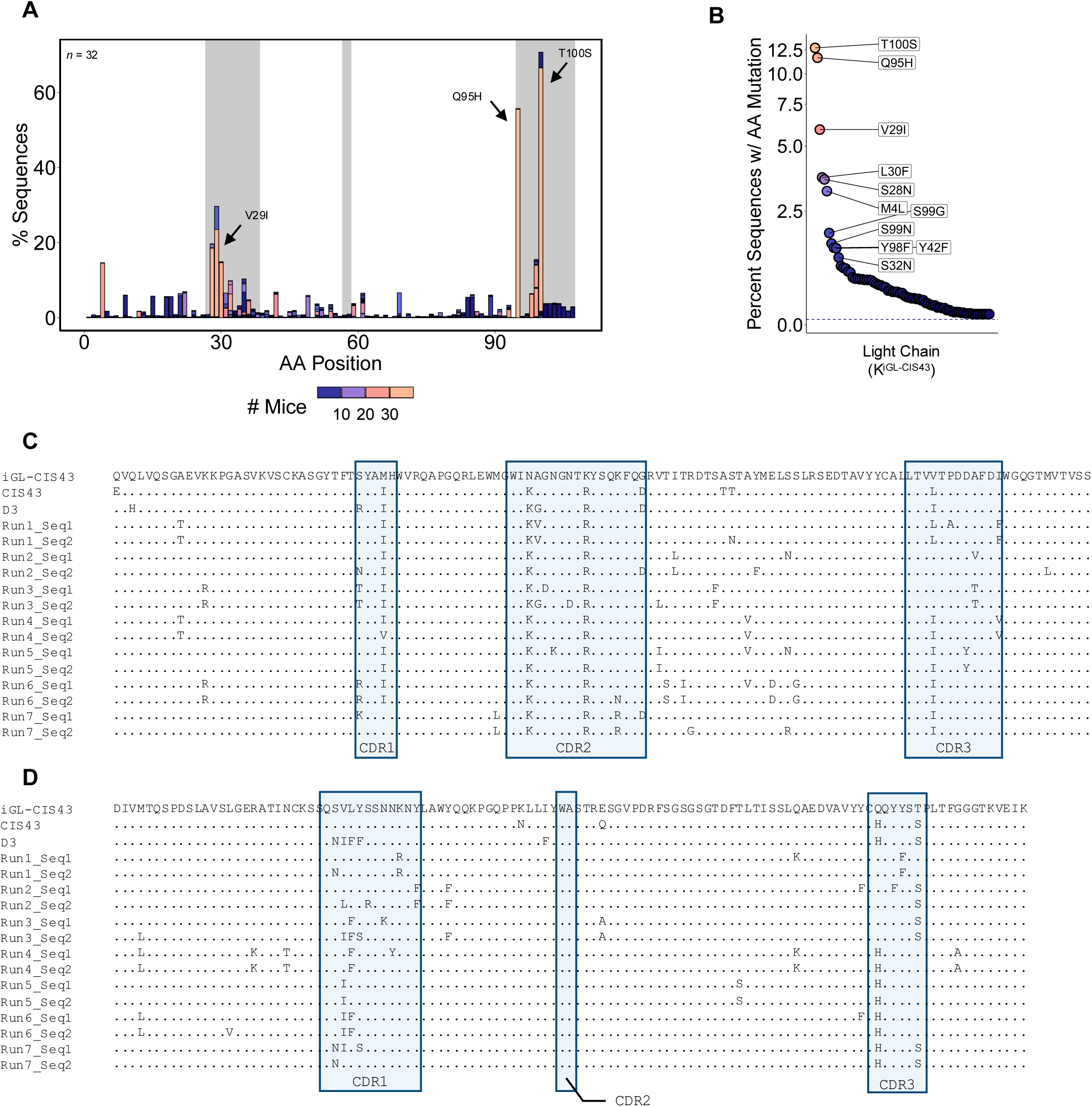
Micro-evolutionarily selected mutations of the light chain in H^iGL-CIS43^K^iGL-CIS43^ expressing B cells following immunization and sequences of isGC-iGL-CIS43 variants. **(A)** Shown are the profile of mutations along the light chain of K^iGL-CIS43^. Data are constructed from 1190 light chain sequences derived from 32 immunized mice as in **Fig. S1**. Mutations are colored to identify amino acid mutations shared across mice. **(B)** Shown are frequency of amino acid mutations following removal of duplicate (clonal) sequences. **(C–D)** Shown are the heavy **(C)** and light **(D)** sequences of the 14 antibodies generated from the isGC using iGL-CIS43 as an input sequence.

**Figure S4:**
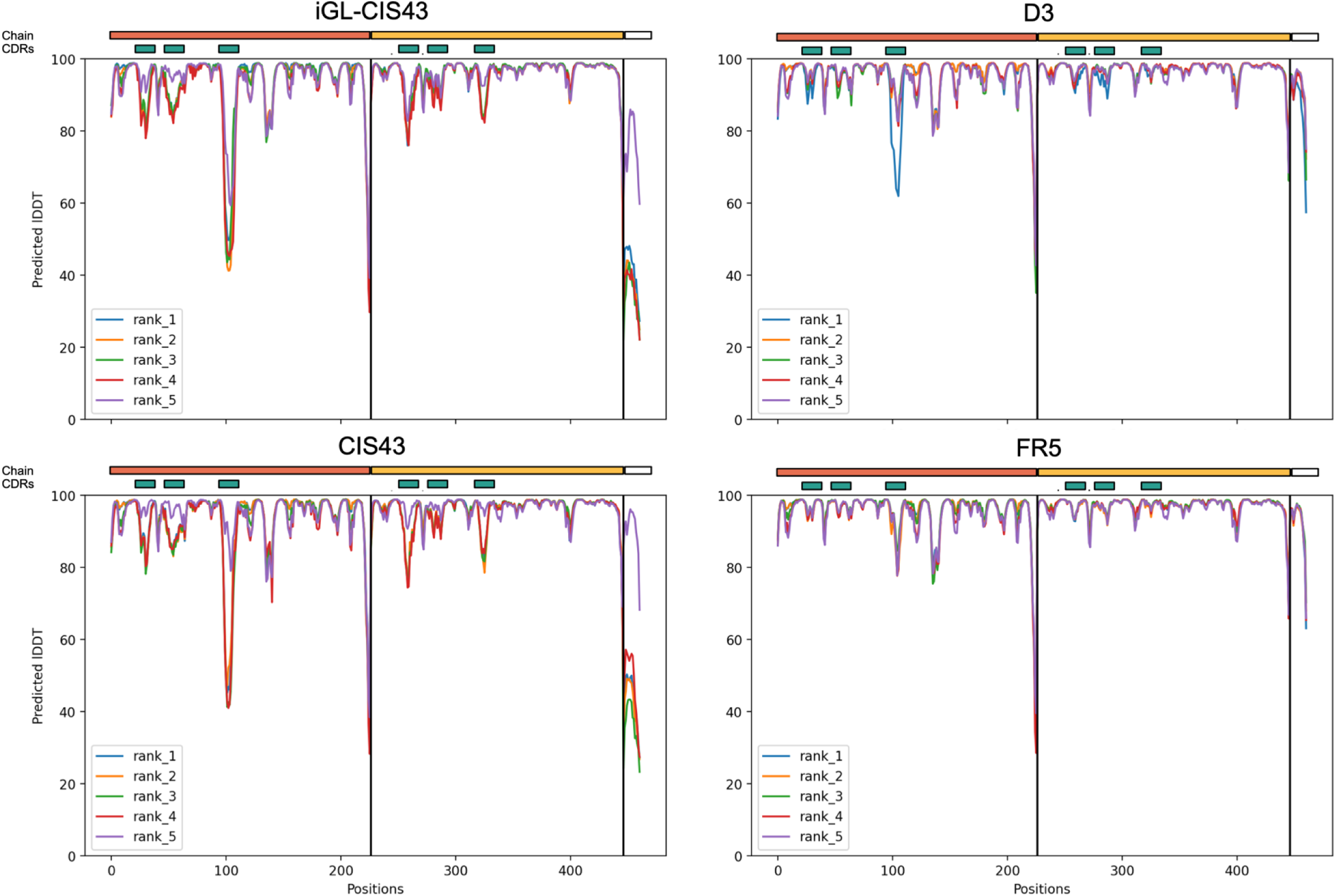
Per residue error plots of iGL-CIS43, CIS43, D3, and FR5 demonstrate the relationship between affinity and AlphaFold pLDDT scores. Shown are pLDDT scores of iGL-CIS43, CIS43, D3, and FR5 complexed with peptide 21. Each plot shows the per residue error at each position along the prediction. Shown are each of the five models predicted by AlphaFold. For each plot, the bar above the plot denotes which chain. Orange denotes the heavy chain, yellow denotes the light chain, and white denotes peptide 21. CDRs are demarcated by teal bars above each plot. Relative affinities are, lowest to highest, iGL-CIS43, CIS43, D3, and FR5.

**Figure S5:**
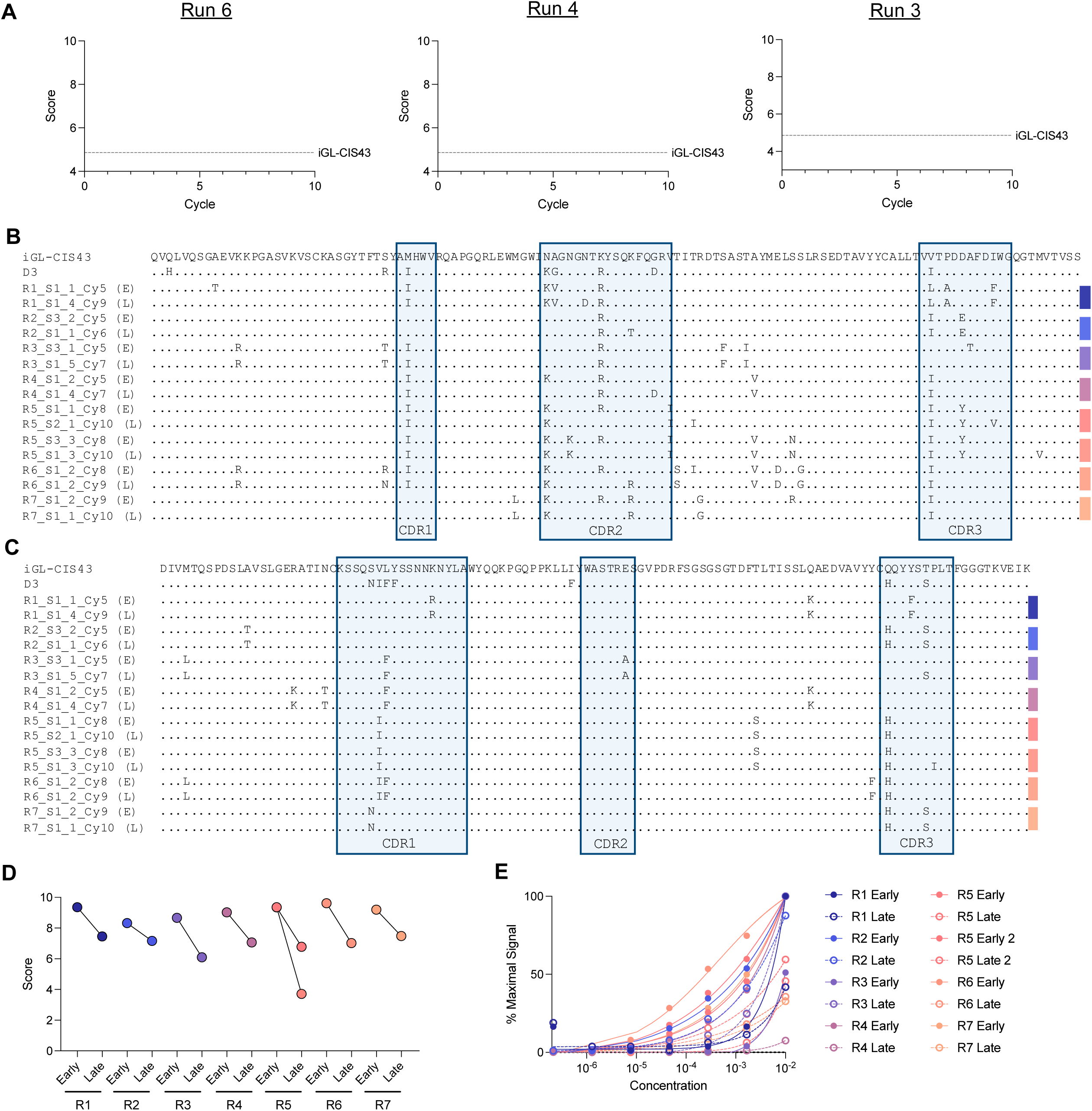
isGC scores correlate with relative affinities of sequences matured from germline. **(A)** Shown are the scores of all sequences produced during an isGC run with most sequences decreasing in score during each cycle. **(B–C)** Shown are the heavy **(B)** and light **(C)** chain sequences from igL-CIS43 isGC runs that were used to understand predictive value of decreases in score through the isGC. Progenitor sequences are termed early (E) or late (L). Colored bars indicate paired sequences. **(D)** Scores of paired sequences in B and C. **(E)** Sequential dilution ELISA results for binding of antibody variants to peptide 21 coated plates. Filled circles denote the early sequences while late sequences are hollow.

**Figure S6:**
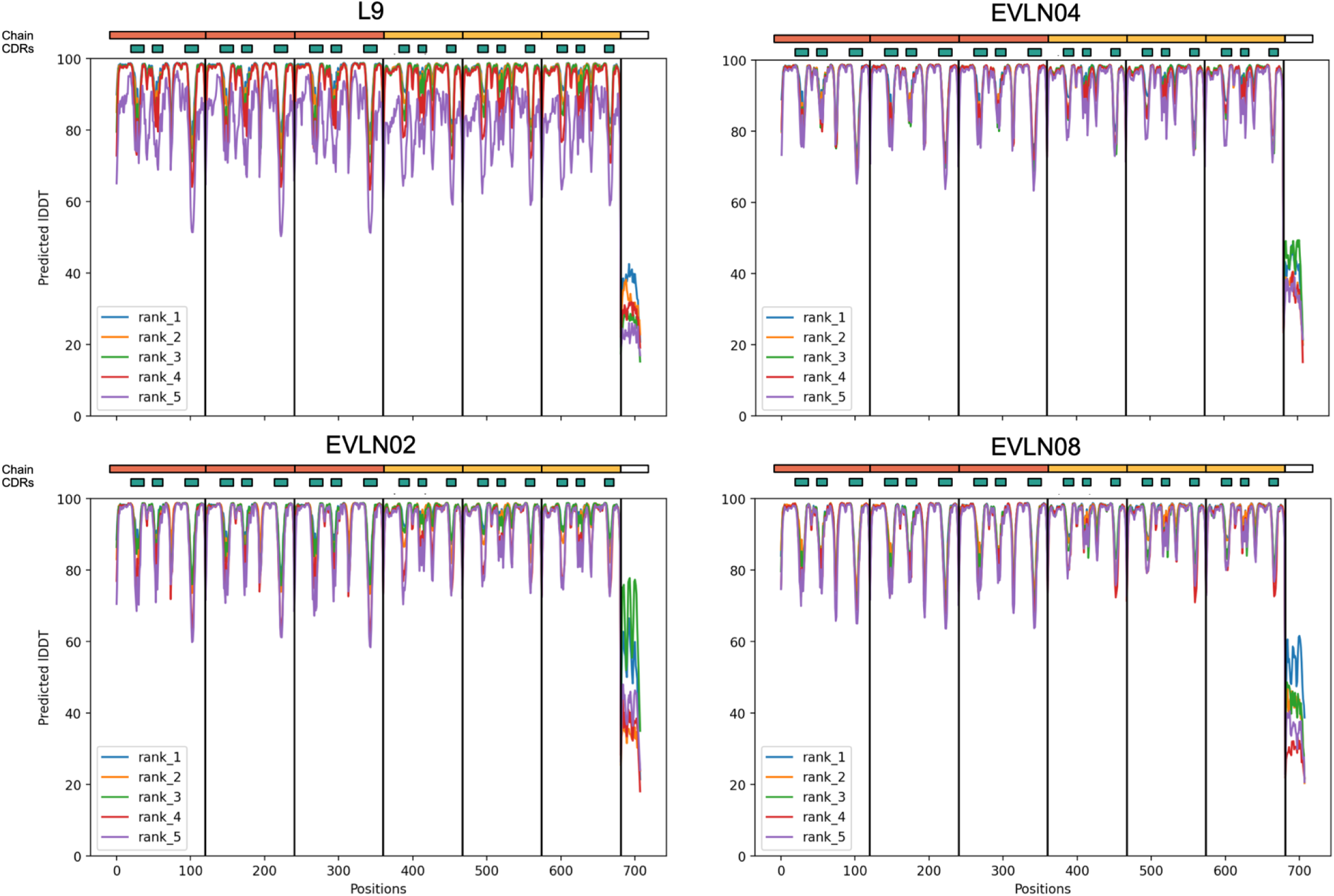
Per residue error plots of L9 and EVLN variants. Shown are pLDDT scores of L9, EVLN02, EVLN04, and EVLN08 complexed with peptide 21-25. Data are presented as in **Fig. S4**.

**Figure S7:**
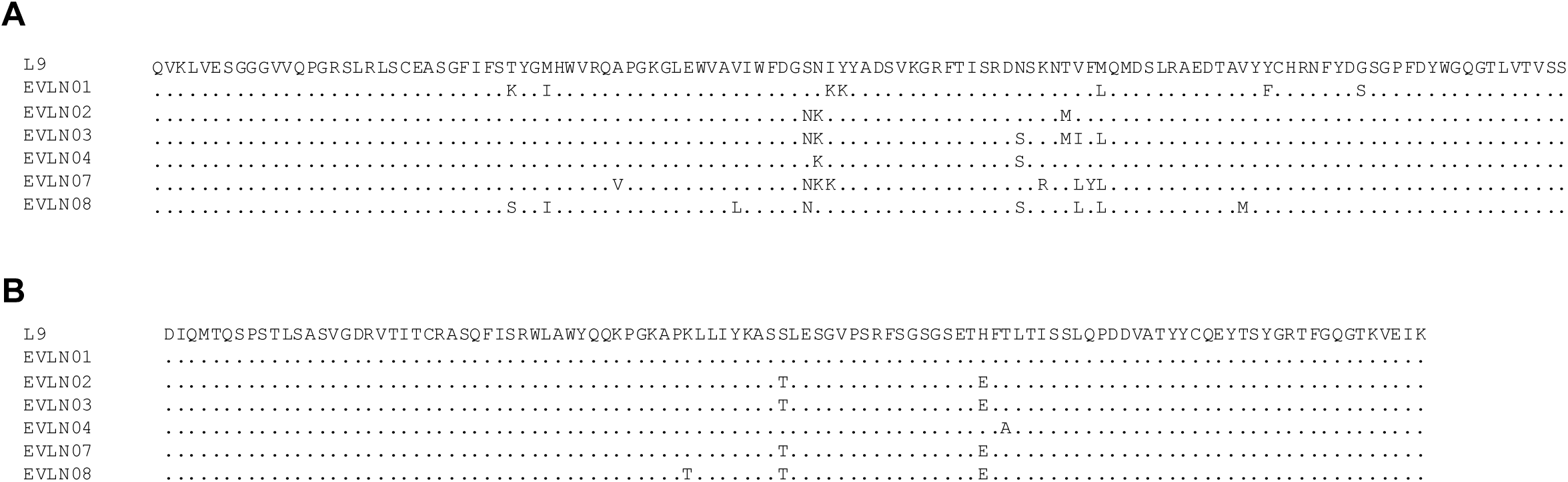
Sequence of L9 and EVLN variants. (A–B) Shown are the heavy chain **(A)** and light chain **(B)** of L9 and the EVLN-series variants. Sequences are depicted as in **Fig. S2**.

